# A conserved protein, BcmA, mediates motility, biofilm formation, and host colonisation in Adherent Invasive *Escherichia coli*

**DOI:** 10.1101/606798

**Authors:** Robert Cogger-Ward, Adam Collins, Denise McLean, Jacob Dehinsilu, Alan Huett

**Author notes:** Corresponding author (A.H.).

## Abstract

Adherent Invasive *Escherichia coli* (AIEC) is a non-diarrhoeagenic intestinal *E. coli* pathotype associated with Crohn’s Disease. AIEC pathogenesis is characterised by biofilm formation, adhesion to and invasion of intestinal epithelial cells, and intracellular replication within epithelial cells and macrophages. Here, we identify and characterise a protein in the prototypical AIEC strain LF82 which is required for efficient biofilm formation and dispersal – LF82_p314. LF82 Δ*LF82_314* have defective swimming and swarming motility, indicating LF82_p314 is important for flagellar-mediated motility, and thus surface colonisation and biofilm dispersal. Flagellar morphology and chemotaxis in liquid appear unaffected by deletion of *LF82_314*, suggesting LF82_p314 does not elicit an effect on flagella biogenesis or environmental sensing. Flagellar motility has been implicated in AIEC virulence, therefore we assessed the role of LF82_p314 in host colonisation using a *Caenorhabditis elegans* model. We found that LF82 Δ*LF82_314* have an impaired ability to colonise the *C. elegans* compared to wild-type LF82. Phylogenetic analysis showed that *LF82_314* is conserved in several major enterobacterial pathogens, and suggests the gene may have been acquired horizontally in several genera. Our data suggests LF82_p314 may be a novel component in the flagellar motility pathway and is a novel determinant of AIEC colonisation. Our findings have potential implications not only for the pathogenesis of Crohn’s Disease, but also for the course of infection in several major bacterial pathogens. We propose a new designation for *LF82_314*, <u>*b*</u>iofilm <u>*c*</u>oupled to <u>*m*</u>otility <u>*A*</u>, or *bcmA*.

**Author summary:** Adherent Invasive *Escherichia coli* (AIEC) are a group of bacteria implicated in the pathogenesis of Crohn’s Disease, a chronic inflammatory bowel disease with no cure. Critical to the process of many bacterial infections is the ability of bacteria to swim towards and colonise the host surface using specialised, propeller-like appendages called flagella. In this paper, we describe a novel protein – LF82_p314 (BcmA) – which is required for efficient flagella-mediated motility and surface colonisation in AIEC. Using a nematode worm (*Caenorhabditis elegans*) infection model, we show that LF82_p314 enables effective colonisation of the *C. elegans* gut, suggesting a role for the protein during human infection. These findings indicate BcmA is significant for initial colonisation of the human gut by AIEC, and therefore the onset of Crohn’s Disease.

## Introduction

Crohn’s Disease (CD) is a chronic and relapsing inflammatory bowel disease presenting with frequent bloody diarrhoea, bowel obstruction, abdominal pain, and extraintestinal manifestations affecting the eyes, skin, joints, and liver (reviewed in [1–3]). CD is a complex syndrome which is understood as an unchecked and inappropriate inflammatory response to intestinal bacteria, potentiated by carriage of one or several of over 180 predisposing immune-related alleles [4–13] and their interaction with environmental risk factors such as smoking [14–16]; consumption of a “western” high-fat, low-fibre diet [17–20]; colonisation by a low-complexity, pro-inflammatory microbiome [21–23]; and carriage of CD-associated pathobionts *Mycobacterium avium* subsp. *paratuberculosis* [24,25] and Adherent Invasive *Escherichia coli* (AIEC) [26–34]. An increasing body of evidence suggests AIEC can act as a key aetiological component of CD. AIEC strains are found present in the ileal mucosa of up to 51.9% CD patients compared to 16.7% healthy controls [26,30,33], and have been shown to induce inflammation and colitis in mice carrying CD-associated *TLR5* deletions [35–37]; mice fed CD-associated “western” diets [19]; and mice with infection-associated intestinal inflammation and microbiome perturbations [37–39]. Indeed, the recent demonstration that AIEC alone can perturb simple microbiomes and instigate inflammation in a *TLR5^−/−^* mouse model [37] raises the possibility that – given a set of predisposing factors – AIEC infection may serve as a first step towards triggering the CD inflammatory cascade.

AIEC pathogenesis is classically characterised by adherence to, invasion of, and replication within intestinal epithelial cells (IECs) and macrophages [40–43]. Despite its significance in CD aetiology, however, the molecular pathogenesis of AIEC infection is comparatively poorly understood. AIEC are thought to use flagella to swim through the mucus layer in the gut [44–46], and secrete the mucolytic Vat-AIEC protease [47] to gain access to the intestinal epithelial surface. AIEC bind the epithelial surface via long polar fimbriae [48] and interactions between type 1 pili and CEACAM6 [40,49–51], a host adhesin over-expressed by CD patient intestinal epithelial cells. Epithelium-associated AIEC may be transcytosed by microfold cells into Peyer’s Patches to be phagocytosed by macrophages, may actively invade IECs, or may alternatively form biofilms on the luminal surface of the gut. Invasion is mediated by microtubule polymerisation and actin recruitment [41], and is thought in part to be facilitated *via* uncharacterised effector delivery in outer-membrane vesicles (OMVs) [52,53], and by a putative oxidoreductase, *ibeA* [54]. However, inhibited OMV release and *ibeA* deletion do not fully abrogate invasion, suggesting other, unknown factors may be involved. The mechanisms of intracellular replication remain to be elucidated, with only one protein – the oxidoreductase *dsbA* – known to be required [55]. In addition to canonical adhesion and invasion traits explored in the first descriptions of AIEC, biofilm formation [56–58], motility [44–46,59], and the ability to utilise short-chain fatty acids (SCFA’s) as carbon sources [59–61] are becoming understood as determinants of AIEC pathogenesis. The discovery of elevated mucosa-associated biofilms in CD patients [29] suggests biofilms may be of specific importance in AIEC pathogenesis, warranting further investigation.

We previously conducted a high-throughput heterologous expression screen to identify putative effectors in the prototypical AIEC strain, LF82 [62]. AIEC-specific putative virulence genes were selected by comparison of the LF82 genome to several pathogenic and commensal *E. coli* reference genomes, and expressed in a HeLa cell line as GFP fusions. Automated microscopy and image analysis of putative effector-GFP fusions expressing HeLa cells allowed identification of protein subcellular localisations and co-localisations. From this screen, we identified a conserved hypothetical protein of unknown function – *LF82_314* – which self-assembles into large filaments (Fig. S1). *LF82_314* is widely conserved, and bioinformatic analysis (Table S1) suggested the gene is co-inherited with components of the General Secretion Pathway (GSP). The GSP is required for secretion of extracellular proteins, including pili (reviewed in [63]). We therefore hypothesised that *LF82_314* may encode either a novel, self-assembling pilin, or an amyloid-like biofilm matrix component. Using established biofilm and motility assays, and a *Caenorhabditis elegans* infection model, we established that LF82_p314 is required for efficient biofilm formation, motility, and host colonisation in LF82. Furthermore, bioinformatic analysis reveals *LF82_314* is conserved in a range of enterobacterial genomes, many of which are human pathogens. Because of the roles *LF82_314* plays in infection, and the potential significance of this novel virulence factor in diverse enterobacterial pathogens, we propose a new designation for *LF82_314*, <u>*b*</u>iofilm <u>*c*</u>oupled to <u>*m*</u>otility <u>*A*</u>, or *bcmA*.

## Results

### LF82_p314 promotes biofilm formation

To identify a role for LF82_p314 in biofilm formation, we created a clean, markerless deletion in *LF82_314*, LF82 Δ*LF82_314*. Using a microtitre plate-based crystal violet assay, we established that LF82 Δ*LF82_314* has a marked biofilm formation defect with incomplete dispersal upon biofilm maturation, when compared to wild-type LF82 (Fig. 1A). Episomal expression of LF82_p314 from p*LF82_314* complemented *LF82_314* deletion. Microscopic analysis of LF82 biofilms formed on glass cover slips revealed LF82 Δ*LF82_314* form patchier, less complete biofilms than wild-type LF82, a defect which can be also complemented by LF82_p314 expression (Fig. 1B). We theorised that the biofilm formation defect may be due to defective initial surface attachment, intercellular adhesion, or altered extracellular matrix architecture. If intercellular adhesion or extracellular matrix formation is altered by *LF82_314* deletion, LF82 Δ*LF82_314* biofilm formation may be complemented in *trans* by co-culture with wild-type LF82. We therefore conducted a *trans*-complementation assay, in which the biofilm formation of wild-type LF82 and LF82 Δ*LF82_314* mixed in a 1:1 ratio was assessed. We found that the LF82:LF82 Δ*LF82_314* mix formed biofilms of intermediate mass when compared to LF82 and LF82 Δ*LF82_314* biofilms (Fig 1C). To characterise the architecture of these mixed biofilms, LF82 and LF82 Δ*LF82_314* strains expressing sGPF2 and mScarlet-I, respectively, were generated for fluorescence microscopy of biofilms (Fig. 1D). LF82-sGFP2 and LF82 Δ*LF82_314*-mScarlet-I biofilms appear similar in extant and structure to those generated by non-fluorescent, parental strains. When mixed in a 1:1 ratio, LF82-sGFP2 and LF82 Δ*LF82_314*-mScarlet-I form biofilms composed of distinct, strain-exclusive islands, suggesting initial attachment and biofilm growth of LF82 and LF82 Δ*LF82_314* are independent of one another. Taken together with Fig. 1C, this demonstrates that the biofilm formation defect observed in LF82 Δ*LF82_314* cannot be complemented in *trans*, suggesting that LF82_p314 is unlikely to have a direct role in intercellular adhesion or biofilm matrix architecture.

**Fig 1.**
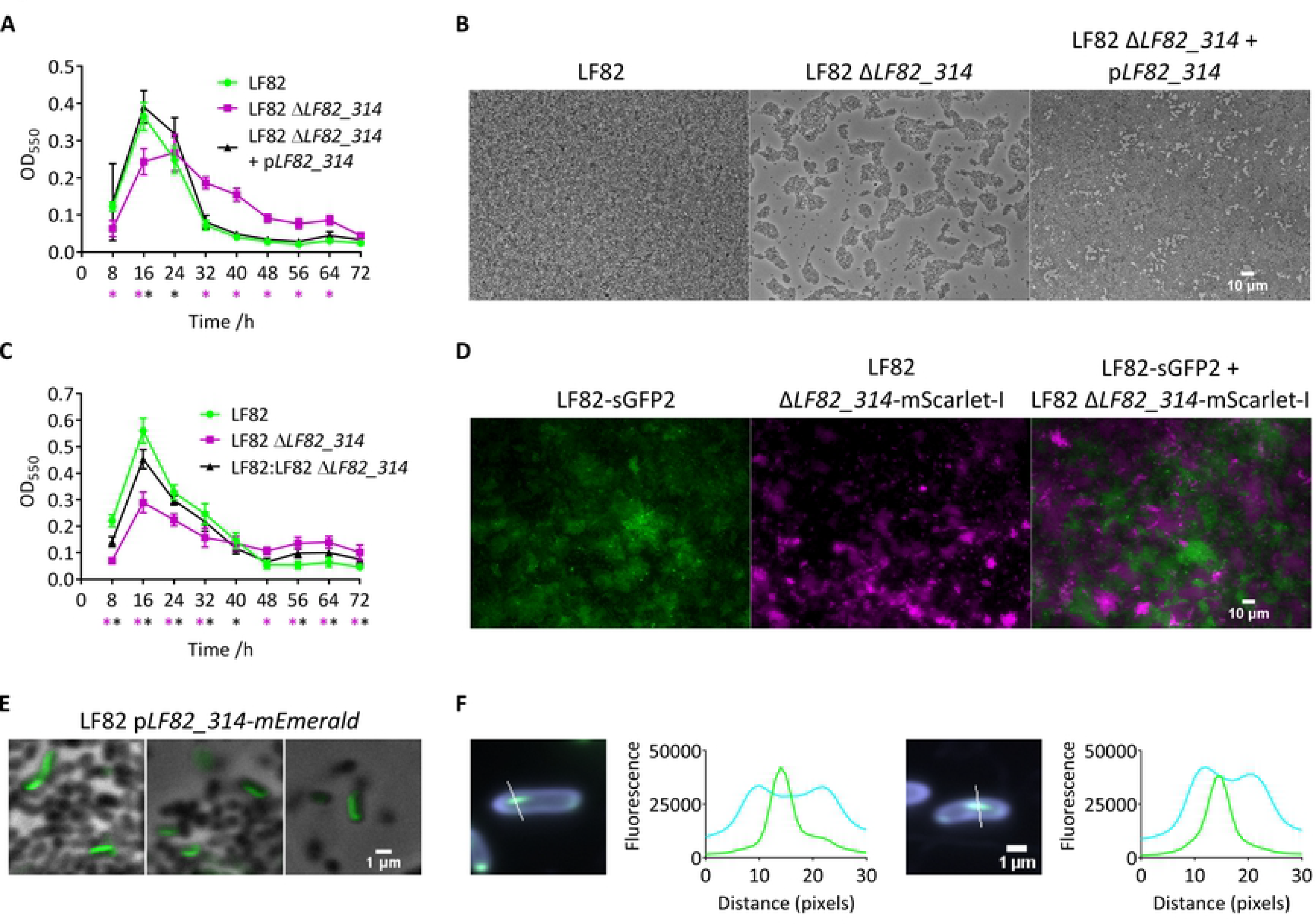
*LF82_314* is required for optimal biofilm formation. (A) LF82 Δ*LF82_314* do not form biofilms as strongly as wild-type LF82, and LF82 Δ*LF82_314* biofilms do not appear to mature and disperse as readily as wild-type biofilms. Wild-type biofilm formation and dispersal behaviour is restored by episomal expression of *LF82_314*. Coloured asterisks represent significant difference between mutant (magenta) and complemented (black) groups, and LF82 (two-way ANOVA with multiple comparisons to LF82; see Table S2 for significance levels). (B) Microscopic analysis of biofilms at 16 h shows LF82 Δ*LF82_314* form patchy, less dense biofilms than wild-type LF82, and that this phenotype can be complemented by *LF82_314* expression. (C) To assess whether LF82_p314 functions in *trans*, we assessed the biofilm formation of wild-type LF82, LF82 Δ*LF82_314*, and a 1:1 mix of LF82:LF82 Δ*LF82_314*. An LF82 Δ*LF82_314* biofilm defect was observed as in (A); however, a 1:1 mix of LF82:LF82 Δ*LF82_314* displayed an intermediate phenotype. Coloured asterisks represent significant difference between mutant (magenta) and mixed (black) cultures, and LF82 (two-way ANOVA with multiple comparisons to LF82; see Table S1 for significance levels). (D) LF82-sGFP2 (green) and LF82 Δ*LF82_314*_mScarlet-I (magenta) form biofilms comparable to non-fluorescent LF82 and LF82 Δ*LF82_314*. A 1:1 co-culture of LF82-sGFP2 and LF82 Δ*LF82_314*_mScarlet-I show that mixed biofilm are composed of strain-exclusive islands. (E) Biofilms in which LF82 express an LF82_p314-mEmerald fusion protein (green) shows LF82_p314 forms cell-associated filaments, as observed in HeLa cells [62]. (F) Fluorescence intensity cross-section analysis of LF82 expressing *LF82_314-mEmerald* with CF633 succinimidyl ester-stained outer membrane proteins shows the extracellular stain and mEmerald fluorescence intensity peaks do not overlap, demonstrating an intracellular localisation for *LF82_314* filaments. White lines represent the 30 pixels of the cross-section. Cyan = CF633; green = LF82_p314-mEmerald.

To define whether LF82_p314 is likely to function as a pilin, adhesin required for initial attachment, or extracellular matrix component, we characterised the cellular localisation of LF82_p314 in biofilms and planktonic cells. Fluorescence microscopy of biofilms formed by LF82 expressing an LF82_p314-mEmerald fusion protein showed LF82_p314 localises as cell-associated filaments – as in HeLa cells (Fig. 1E) – which align with the long axis of the bacterium. To assess the subcellular localisation of the LF82_p314 filaments, we stained live, planktonic LF82 p*LF82_314-mEmerald* with an amine-reactive succinimidyl ester dye conjugate (CF™ 633, Sigma Aldrich) to define the outer membrane, fixed the dyed cells, and them imaged by fluorescence microscopy. Pearson correlation analysis of pixel intensities was performed using CellProfiler and demonstrated a very weak correlation between CF633 and LF82_p314-mEmerald fluorescence (Pearson correlation coefficient, mean *r* = 0.135 ± 0.032 (95% CI)). Furthermore, cross-sectional analysis of fluorescence intensity (Fig. 1F) shows two peaks of CF633 intensity, representing the cell membranes, and one peak of LF82_p314-mEmerald intensity between these peaks, suggesting that LF82_p314 filaments localise intracellularly. We also note that in cells imaged 1 hour post induction, LF82_p314 filaments are shorter than in biofilms imaged at 16 h, and localise near the cell pole, suggesting interactions with intracellular, pole-localised proteins. LF82_p314 is therefore unlikely to be a pilin or extracellular matrix component, and the biofilm defect observed in LF82 Δ*LF82_314* is not due to aberrant pilin-mediated attachment or extracellular matrix architecture.

### LF82_p314 modulates flagella-mediated motility *via* an uncharacterised mechanism

In the absence of evidence for an adhesin or extracellular matrix function for LF82_p314, we reasoned that a motility defect might confer a surface colonisation defect, manifesting in an apparent biofilm formation defect, as has been shown elsewhere [64]. Accordingly, we used established soft agar motility assay methods to analyse the swimming and swarming behaviour of LF82, LF82 Δ*LF82_314*, and LF82 Δ*LF82_314* p*LF82_314*. We found that LF82 Δ*LF82_314* has notable swimming (Fig. 2A) and swarming (Fig. 2B) defects (One-way ANOVA with multiple comparisons to wild-type LF82; swim, LF82 vs LF28 Δ*LF82_314*, p = 0.0004; swarm, LF82 vs LF28 Δ*LF82_314*, p = 0.0001) when compared to wild-type LF82 at 10 and 24h post-inoculation, respectively. No significant difference was observed between LF82 and the *LF82_314*-expressing strain, LF82 Δ*LF82_314* + p*LF82_314*, demonstrating these defects are fully complemented by *LF82_314* expression. We noted that at 24h post-inoculation, both LF82 and LF82 Δ*LF82_314* on swimming plates had reached the edge of the plate; however, these plates lack the characteristic chemotactic rings observed on wild-type LF82 and LF82 Δ*LF82_314* + p*LF82_314* plates (Fig. 2A), and also often showed swarming behaviour in the centre. We therefore theorised that the motility and/or chemotaxis systems may be defective in LF82 Δ*LF82_314*, leading to a slower rate of swimming, and/or an inappropriate response to wetness conditions. To test the chemotactic response of LF82 Δ*LF82_314*, we conducted a simple capillary-based chemotaxis assay using media with or without glucose as a chemoattractant (Fig. 2C). We found both LF82 and LF82 Δ*LF82_314* are more enriched in capillaries containing glucose than without, and no statistically significant difference was observed, suggesting chemotaxis is intact in LF82 Δ*LF82_314*. We also assessed whether the number per cell or morphology of flagella was affected by deletion of *LF82_314*, using Kodaka staining [65] and transmission electron microscopy (TEM). Kodaka staining (Fig. 2D) confirmed the presence of flagella on both wild-type LF82 and LF82 Δ*LF82_314*. Negative-stain TEM demonstrated no gross morphological differences in flagella between wild-type LF82 and LF82 Δ*LF82_314* flagella (Fig. 2E). Flagella counts from 30 TEM micrographs (Fig. 2F) revealed no difference between the numbers of flagella per flagellated cell. These data demonstrate *LF82_314* is required for efficient flagella-mediated motility; however, gross behavioural and morphological traits such as in-liquid chemotaxis and flagella biosynthesis are intact in LF82 Δ*LF82_314*, suggesting LF82_p314 elicits its effect via a more subtle, uncharacterised mechanism.

**Fig. 2.**
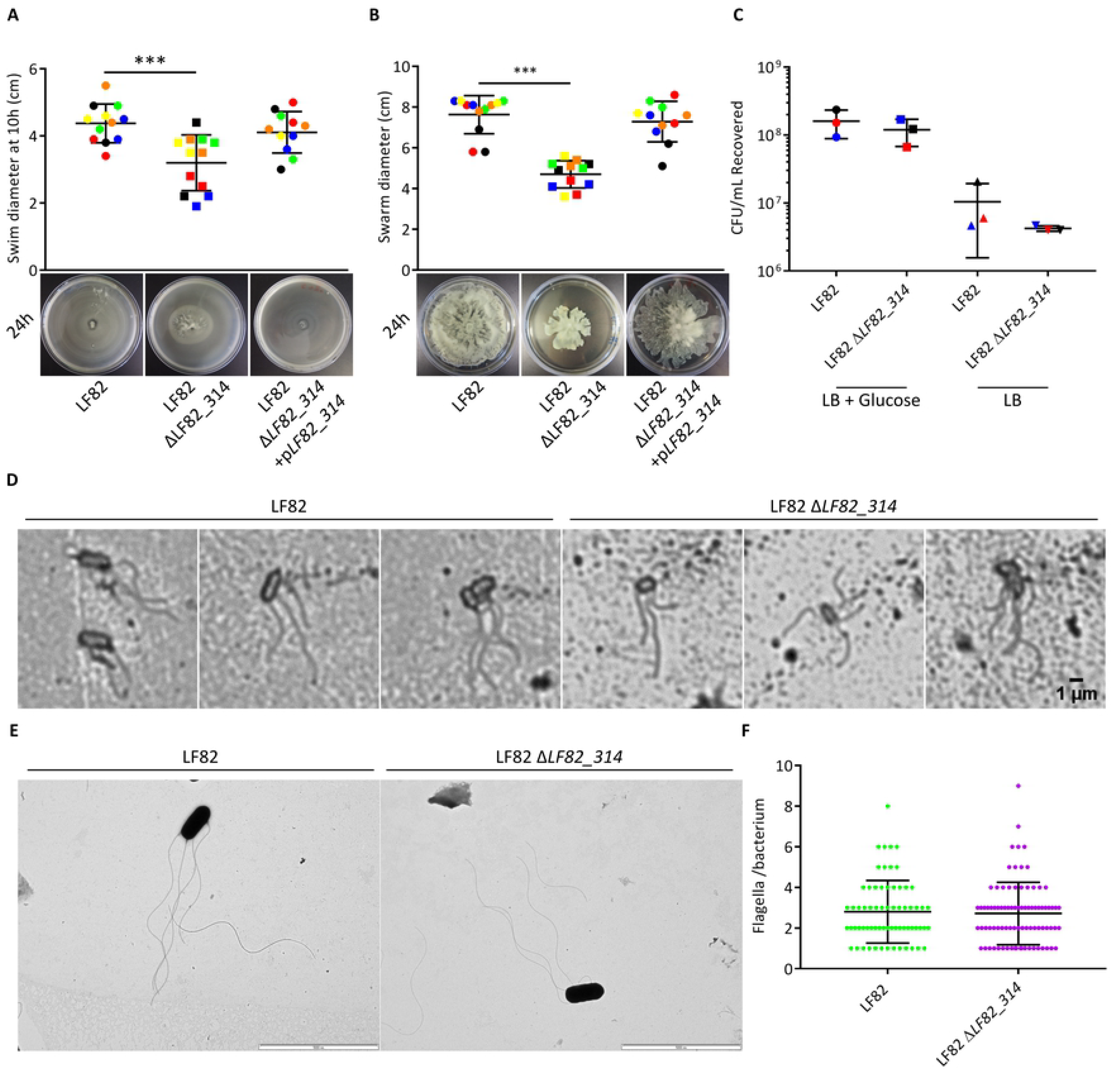
*LF82_314* promotes flagella-mediated motility *via* an uncharacterised mechanism. LF82 Δ*LF82_314* has notable defects in (A) swimming and (B) swarming motility (One-way ANOVA, *** = p ≤ 0.001), which are complemented by *LF82_314* expression. Each dot represents one technical replicate, or plate; separate colours represent biological replicates. Swim plates were measured at 10 h post-inoculation, and swarm plates at 24h. Motility plates imaged at 24 h post-inoculation show LF82 Δ*LF82_314* have atypical swimming motility lacking chemotactic rings observed on wild-type plates, and the swarming defect. (C) A capillary-based chemotaxis assay demonstrated increased recovery of both LF82 and LF82 Δ*LF82_314* CFU from media supplemented with glucose compared to LB alone, with no significant difference between strains in CFU recovered in either condition. Each dot represents one biological replicate. Bright-field microscopy of Kodaka stained (D) and negative stain TEM (E) of LF82 and LF82 Δ*LF82_314* demonstrates no differences in flagella morphology, and flagella counts from 30 TEM micrographs (F) show no difference in flagella numbers per flagellated cell, suggesting flagella biosynthesis is intact in both strains. TEM scale bar represents 5 µm.

### LF82_p314 is required for optimal *C. elegans* gut colonisation

Non-motile AIEC have significantly reduced virulence in *in vivo* models [45,46,66], and host-adapted AIEC are hyper-motile [59], suggesting flagella motility is critical in AIEC virulence. We therefore assayed the *in vivo* virulence of LF82 and LF82 Δ*LF82_314* using an established *C. elegans* survival assay [66]. The *C. elegans* food source strain *E. coli* OP50 was used as a negative control. We found LF82 was capable of “slow killing” *C. elegans*, and that deletion of *LF82_314* does not improve or abrogate the survival of infected *C. elegans* (Fig. 3A), suggesting *LF82_314* does not directly contribute to *C. elegans* killing by AIEC in this model. We noted, however, that worms fed wild-type LF82 consistently begin to die 1-2 days before those fed LF82 Δ*LF82_314*, and reasoned that this may be due to less efficient colonisation of the *C. elegans* gut by the less motile LF82 Δ*LF82_314*, prolonging the time required to fully establish infection. We therefore chose to assess the number of bacteria stably colonising the *C. elegans* gut at daily intervals. In worms fed on lawns containing exclusively LF82 or LF82 Δ*LF82_314*, we found no statistically significant deviation between wild-type and mutant CFU recovered per worm gut (Fig. 3B), although mean LF82 CFU per worm gut was higher at 3 and 4 days post-infection (d.p.i.).

**Fig. 3.**
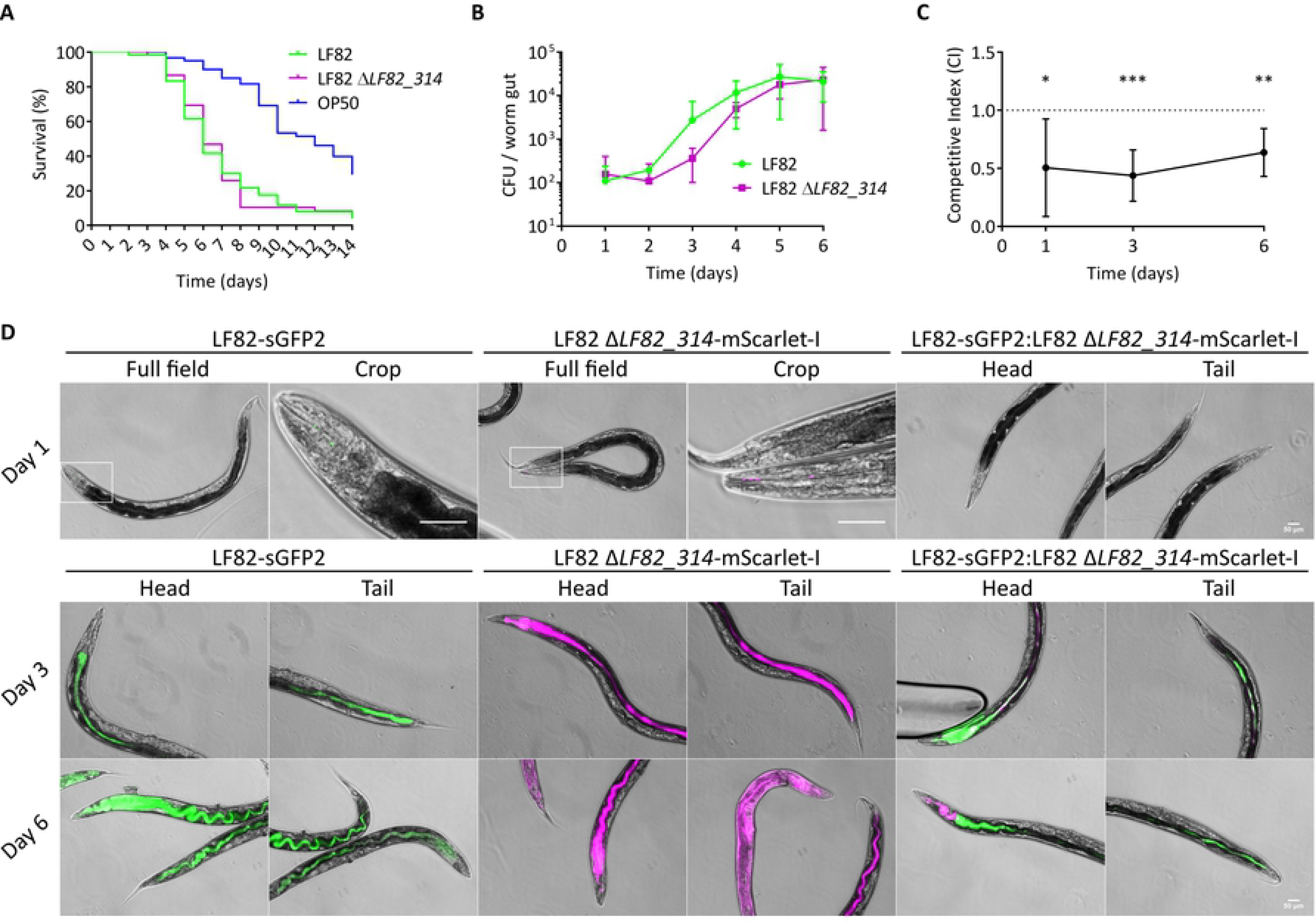
*LF82_314* promotes gut colonisation in *Caenorhabditis elegans*. (A) Survival of *C. elegans* SS104 is significantly decreased when cultivated on LF82 or LF82 Δ*LF82_314*, compared to OP50, however no significant difference was noted between survival on LF82 or LF82 Δ*LF82_314*. (B) No significant difference in stable colonisation of the *C. elegans* gut by LF82 or LF82 Δ*LF82_314* was found when worm were fed on each strain exclusively. (C) A competition assay demonstrated LF82 Δ*LF82_314* has a competitive colonisation disadvantage compared to LF82 throughout the course of infection (one-tailed Wilcoxon match-pairs signed rank test, * = p ≤ 0.05, ** = p ≤ 0.01, *** = p ≤ 0.001. CI’s were compared to an ideal “no-disadvantage” CI = 1). CI ratios below 1 represent a competitive disadvantage. Data was pooled from 3 independent experiments. Error bars show 95% confidence intervals. (D) Fluorescence microscopy of *C. elegans* fed LF82-sGFP2 and LF82 Δ*LF82_314*-mScarlet-I alone demonstrates both LF82 and LF82 Δ*LF82_314* are capable of establishing gut and pseudocoelomic infections; however, when fed to worms in a 1:1 ratio, LF82-sGFP2 appears to outcompete LF82 Δ*LF82_314*-mScarlet-I, mirroring Fig. 3C. These data suggest that *LF82_314* is required for efficient host colonisation. Scale bars represent 50 µm.

Reasoning that an assay in which *C. elegans* are continuously fed up to 10^11^ CFU/mL of one bacterial strain may not represent a realistic infection scenario, and that this could mask a colonisation defect, we conducted competition assays to detect whether *LF82_314* deletion impacts fitness against wild-type LF82. LF82 and LF82 Δ*LF82_314* carrying the Kanamycin-resistant pBAD18 (LF82-Kan^R^ and LF82 Δ*LF82_314*-Kan^R^) or Chloramphenicol-resistant pBAD33 (LF82-Cm^R^ and LF82 Δ*LF82_314*-Cm^R^) were used to allow differential selection of CFU recovered from *C. elegans*. Worms were fed on 1:1 mixes of LF82-Kan^R^:LF82 Δ*LF82_314*-Cm^R^ or LF82-Cm^R^:LF82 Δ*LF82_314*-Kan^R^ as above. The Competitive Indices (CI) for LF82 Δ*LF82_314* in this assay (Fig. 3C) are significantly below a “no-disadvantage” CI ratio of 1 throughout infection (one-tailed Wilcoxon match-pairs signed rank test, 1 d.p.i., p = 0.0195; day 3 d.p.i., p = 0.0004; day 6 d.p.i., p = 0.0011), showing LF82 Δ*LF82_314* has a gut colonisation disadvantage to wild-type LF82. To visualise the infection process, LF82-sGFP2 and LF82 Δ*LF82_314*-mScarlet-I were used to infect worms either alone, or mixed in a 1:1 ratio as above. Fluorescence microscopy of infected worms (Fig. 3D) shows that at 1 d.p.i., both LF82-sGFP2 and LF82 Δ*LF82_314*-mScarlet-I colonise the worm mouth. Fluorescence in worms fed a LF82-sGFP2:LF82 Δ*LF82_314*-mScarlet-I mix was below background levels. At 3 d.p.i. and 6 d.p.i, LF82-sGFP2 and LF82 Δ*LF82_314*-mScarlet-I successfully colonise the head and gut of worm in both mono- and co-feeding conditions, with penetration into the pseudocoelom at 6 d.p.i.; however, in concordance with Fig. 3C, LF82-sGFP2 appears to outcompete LF82 Δ*LF82_314*-mScarlet-I when co-fed to *C. elegans*. Taken together, our data suggests that although deletion of *LF82_314* does not attenuate “slow killing” of *C. elegans* in mono-feeding conditions, *LF82_314* is required for efficient colonisation of the *C. elegans* gut.

### *LF82_314* is a widely-conserved, and may be horizontally transmissible

*LF82_314* is encoded by 468 DNA bases, annotated as encoding the 155 residue protein LF82_p314 [67]. All results returned by blastp and JACKHMMER searches designate *LF82_314* as a “hypothetical protein,” “conserved hypothetical protein,” “MULTISPECIES: hypothetical protein,” “uncharacterised protein,” or “conserved uncharacterised protein”. *LF82_314* is located proximal to a tRNA site (*asnV*) in a region of the LF82 genome (Fig. 4A) which encodes predicted transposases (*LF82_309* and *yhhI*), integrases (*LF82_309* and *LF82_311*), a toxin-antitoxin addiction module (*LF82_312* and *LF82_313*), a transcription factor (*LF82_774*), an endonuclease (*LF82_317*), and a helicase (*LF82_318*). The putative components of this genome neighbourhood and its proximity to a common transposable element insertion site (tRNA) led us to theorise that *LF82_314* may be encoded on an active or former mobile genetic element (MGE). MGEs are significant sources of horizontally acquired virulence factors, notable examples of which include the Shiga toxin – which is transmissible among *E. coli* strains by the *stx* bacteriophage, generating highly virulent Shiga Toxin-producing *E. coli* (STEC; reviewed in [68]) – and the *Salmonella* Typhi pathogenicity island, SPI-7 – a mosaic of conjugative elements and temperate bacteriophage insertions which encodes genes for Vi capsule synthesis and the Type III Secretion System effector, *sopE* [69,70].

**Fig. 4.**
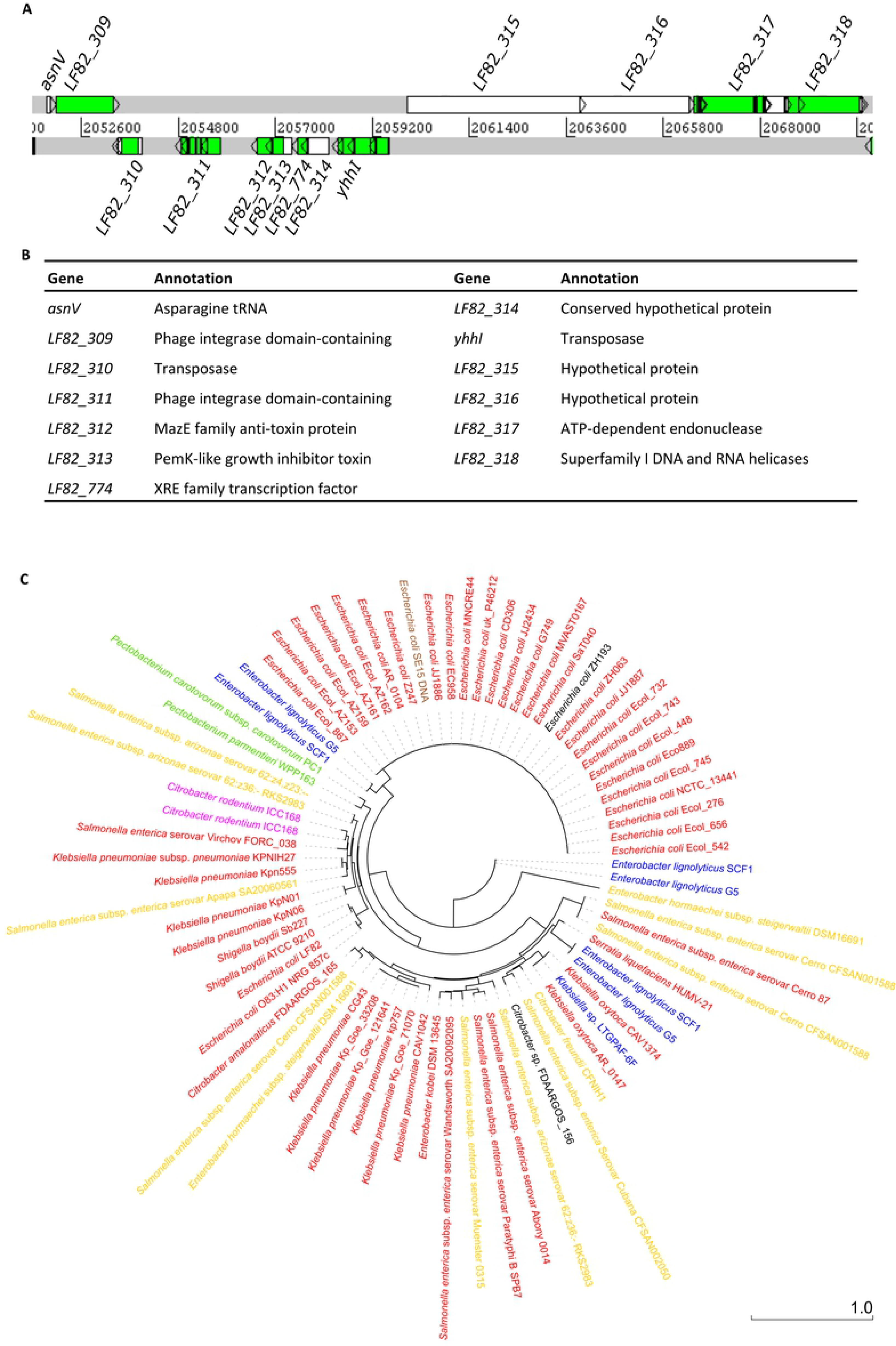
*LF82_314* is conserved among enterobacterial pathogens. *LF82_314* is encoded in a region of the LF82 genome (A) containing several ORFs with predicted transposase and integrase functions, and a toxin-antitoxin addiction module (B), suggesting the region may represent an MGE. (C) A ML tree of 77 *LF82_314* homologues from 68 strains reveals that *LF82_314* is conserved in a wide variety of pathogenic enterobacteria, and that *LF82_314* homologue-derived phylogenies do not recapitulate expected phylogenetic relationships (see Fig. S2). Of note is the large clade *E. coli* strains, which represents members of the clonal, UTI-associated ExPEC, ST131. Red = human pathogen; gold = human and animal or zoonotic pathogen; green = plant pathogen; brown = commensal; blue = environmental; black = insufficient data. Scale bar represents number of substitutions per site.

We therefore sought to assess the distribution of *LF82_314* homologues in related phyla. We harvested the top 100 DNA sequences of *LF82_314* homologues returned by blastn discontiguous megablast (Table S3), and curated this list to remove strains for which 16S rRNA sequences were not readily available. This produced a list of 77 *LF82_314* homologues encoded in 68 enterobacterial genomes. Many of the strains returned by our search strategy are human pathogens (see Fig 4C). Of note are several *E. coli* strains which belong to an emergent clonal, pandemic urinary tract infection (UTI)-associated Extraintestinal Pathogenic *E. coli* (ExPEC) clade, ST131 [71]. Interestingly, blast search strategies excluding the order enterobacteriales did not return any significant results, suggesting *LF82_314* homologues are restricted to this order. We obtained 16S rRNA sequences from the 68 selected strains from SILVA, and used these to build a Maximum Likelihood phylogenetic tree. Comparison of this 16S rRNA tree (Fig. S2) with a Maximum Likelihood tree created from *LF82_314* homologues showed marked differences (Figure 4C). For example, in the *LF82_314* homologue tree, *Salmonella* sp. have a fragmented phylogeny rather than clustering as a distinct phylogenetic group as in the 16S rRNA tree. Similarly, *Shigella boydii* strains, and AIEC LF82 and NRG 857c, cluster together away from the *E. coli* ST131 clade in the *LF82_314* tree, however in the 16S rRNA tree one *E. coli* group is formed. These data suggest that *LF82_314* may have been introduced into these strains horizontally, raising the possibility that in some conditions, *LF82_314* may become a transmissible virulence factor.

## Discussion

Flagella-mediated motility is a critical virulence factor in a wide variety of gram-negative bacterial pathogens. In AIEC LF82 and the closely related strain NRG 857c, flagella motility has been shown to be required for host colonisation, cell invasion, and persistence in *in vivo* models [45,46,59,66]. Flagellar biosynthesis in AIEC is regulated in a canonical fashion by the master flagellar regulators, *flhCD* and *fliA* [44,72]. The *E. coli* quorum sensing system, QseBC, is involved in regulation of *flhCD* function [73], and a novel transcription factor, NrdR, has been implicated in flagellar biosynthesis and regulation of chemotaxis gene expression [45]. The AIEC LF82 genome encodes a full complement of chemotaxis genes [67], and typical chemotactic responses have been observed in LF82 in the literature and this study, suggesting environmental sensing and motile responses are conserved.

We describe in this study a conserved hypothetical gene – *LF82_314* – which has novel functions in biofilm formation and host colonisation, which are mediated by a role in flagellar motility. Although we initially theorised LF82_p314 may function as a self-assembling pilin or biofilm extracellular matrix protein, we have found that LF82_p314 localises within bacterial cells, and is therefore likely a cytoplasmic or periplasmic protein. We have demonstrated that *LF82_314* is required for efficient swimming and swarming in soft agar motility assays; we note, however, in a soft agar swimming assay, that both LF82 and LF82 Δ*LF82_314* reach the edge of the plate at 24h post-inoculation. When considered with the defect observed at 8h post-inoculation, this observation suggests that LF82 Δ*LF82_314* swims and swarms more slowly than wild-type LF82, a phenotype which may be mediated by defective chemotaxis or flagella. LF82 Δ*LF82_314* swim plates lack chemotactic rings, suggesting aberrant chemotaxis may be responsible for the observed defect. However, the in-liquid chemotactic response to glucose, and flagella biosynthesis, are indistinguishable from wild-type LF82 in LF82 Δ*LF82_314*, suggesting neither gross morphological differences nor defective chemotactic signalling can account for the observed motility defect. LF82_p314 must therefore elicit a more subtle effect which nevertheless manifests as a notable motility defect in soft agar. Currently no model of LF82_p314 function exists. We hypothesise that LF82_p314 may be involved in modulating bacterial velocity in high-viscosity environments, or may be involved in surface sensing and transitioning in-liquid motility to surface-associated motility and adhesion. Further study towards a molecular understanding of LF82_p314’s function – including mapping LF82_p314 protein interactions, and studying the effects of LF82_p314 on the AIEC transcriptome – are ongoing in our laboratory.

We demonstrate that *LF82_314* is required for efficient biofilm formation in LF82. Biofilms play a role in Crohn’s Disease pathology [29], and infection-associated biofilms are often sources of persistence, antibiotic resistance, and tolerance [74,75]. Antibiotic therapy is routinely used as an intervention in CD, and is known to temporarily ameliorate symptoms in the majority of patients [76]. However, relapse during treatment is common and reportedly universal [77] when treatment is halted, suggesting inflammation in relapsing CD may be due to outgrowth of antibiotic resistant or surviving, tolerant bacteria, such as those in mucosa-associated AIEC biofilms. LF82_p314 may therefore be of some interest as a potential anti-virulence target which might potentiate more successful antibiotic treatment in CD, by breaking down or inhibiting formation of drug-tolerant AIEC biofilms.

Of particular significance, our work demonstrates *LF82_314* is required for effective colonisation of the gut in a *C. elegans* infection model. We did not observe decreased virulence or colonisation by LF82 Δ*LF82_314* when worms were fed on one strain exclusively, but were able to detect a clear defect when LF82 Δ*LF82_314* was in competition with wild-type LF82. However, this does not imply that *LF82_314* has only a marginal effect on LF82 gut colonisation. In the assay we have adapted from [66], worms are constantly fed on plates prepared with bacterial concentrations between 10^10^ and 10^11^ CFU per ml of culture. In such mono-feeding experiments, it is likely that bacteria at this density saturate the worm gut, bringing equal numbers of wild-type and mutant bacterial cells in contact with the gut surface, thus masking colonisation defects. Indeed, the colonisation defect observed in the competition assay, which still saturates the gut with bacteria, suggests that in more biologically relevant scenarios – such as a substantially reduced total infectious dose of LF82 competing against an established microbiome – LF82 Δ*LF82_314* may have a marked colonisation defect. Further study to establish the role of *LF82_314* in AIEC colonisation in complex polymicrobial contexts is required to test this hypothesis; however, our data provides strong evidence to suggest that LF82_p314-mediated motility plays an important role in host colonisation.

Finally, we report that *LF82_314* is widely distributed throughout the Enterobacteriaceae, including several significant human pathogens, and that the gene appears to be laterally inherited. Our analysis was limited to the top 100 blastn results; however, the wide distribution of closely related *LF82_314* homologues presented in our analysis suggests that this novel virulence factor is likely to be present in an even greater range of enterobacterial pathogens. Although our analysis does not show that the putative *LF82_314* mobile genetic element can be mobilised in the strains we have analysed, it is conceivable that this element may be transmissible from a strain not included in our analysis. This is of particular interest in the context of some of the strains we analysed, such as those belonging to the *E. coli* ST131 clade. ST131 is a clade of ExPEC associated with antibiotic-resistant recurrent UTIs, which was first identified in 2008 [71]. Among the pathogenic characteristics of ST131 are increased biofilm formation and adhesion to epithelial cells, both processes which require flagella motility, and which may be potentiated by LF82_p314. It is thought that many UTIs are seeded from a gut reservoir and colonisation of both epithelial surfaces occurs via similar mechanisms [78]. An ST131 *LF82_314* homologue may play a role both in establishment of a gut niche, as well as subsequent infection of the urinary epithelium. The presence of *LF82_314* in the genomes of numerous strains representing an emergent pathogen raises the possibility that acquisition of *LF82_314* may have been an important step in becoming such a successful pathogen.

Further work is required to understand the molecular function of LF82_p314; to assess its significance in higher-complexity infection systems; and to characterise fully its distribution, and whether this novel virulence factor is transmissible. What is clear is *LF82_314* is a novel player in flagellar-mediated motility with significance for host colonisation and biofilm formation, and is conserved in a range of important human pathogens. We therefore suggest a new designation for *LF82_314* and its homologues – *bcmA* (<u>*b*</u>iofilm <u>*c*</u>oupled to <u>*m*</u>otility <u>*A*</u>) – to facilitate future work without diverse nomenclature confusing the literature.

## Materials and Methods

### Strains and media

*E. coli* LF82, XL-1, and OP50, were grown in Lysogeny Broth (LB) or on LB agar with supplements, antibiotics, and agitation as appropriate, and incubated at 37°C unless otherwise stated. *E. coli* S17-1 carrying pMRE-Tn7-XXX plasmids were maintained at 25°C. *C. elegans* SS104 [*glp-4(bn2)I.*] obtained from the *Caenorhabditis* Genetics Centre were cultured as in [79]. Strains used in this study are listed in Table 1.

**Table 1.**

| Strain | Genotype / Description | Source / Reference |
| --- | --- | --- |
| *Caenorhabditis elegans* SS104 | [*glp-4(bn2)I.*] | Caenorhabditis Genetics Centre |
| *E. coli* LF82 | Prototypical AIEC type strain. Genome sequenced. | Arlette Darfeuille-Michaud, Université Clermont Auvergne |
| *E. coli* LF82 ΔLF82_314 | LF82 with 428 bases of the LF82_314 CDS deleted | This study |
| *E. coli* LF82 ΔLF82_314 pLF82_314 | *E. coli* LF82 ΔLF82_314 with an LF82_314-3xFLAG construct on pBAD18 | This study |
| *E. coli* LF82-sGFP2 | LF82 with chromosomally inserted sGFP2 | This study |
| *E. coli* LF82 ΔLF82_p314-mScarlet-I | LF82 ΔLF82_314 with chromosomally inserted mScarlet-I | This study |
| *E. coli* LF82-Kan<sup>R</sup> | LF82 carrying pBAD18 | This study |
| *E. coli* LF82-Cm<sup>R</sup> | LF82 carrying pBAD33 | This study |
| *E. coli* LF82 ΔLF82_314-Kan<sup>R</sup> | LF82 ΔLF82_314 carrying pBAD18 | This study |
| *E. coli* LF82 ΔLF82_314-Cm<sup>R</sup> | LF82 ΔLF82_314 carrying pBAD33 | This study |
| *E. coli* OP50 | Uracil auxotrophic *C. elegans* food source. Genome sequenced. Tet<sup>R</sup> | Steve Atkinson, University of Nottingham |
| *E. coli* S17-1 $\lambda$ pir | Strain for conjugation of pMre-Tn7-XXX plasmids into LF82 strains | AddGene, (81) |
| *E. coli* XL-1 blue | Common cloning strain |  |

Strains used in this study. Cm<sup>R</sup> = Chloramphenicol resistant; Kan<sup>R</sup> = Kanamycin resistant; Tet<sup>R</sup> = Tetracycline resistant.

### Genetic manipulation

Genes of interest were amplified by PCR amplification using Phusion-HF DNA Polymerase (NEB). Sequences were inserted into pBAD18 or pBAD33 by restriction digest using EcoRI, KpnI, and XbaI restriction enzymes (NEB), and ligation using T4 DNA ligase (NEB). LF82 Δ*LF82_314* was generated from LF82 wild-type using the CRISPR-Cas9-based no-SCAR strategy [80]. Deletion was confirmed by Sanger sequencing, and strains were fully validated by whole genome Illumina sequencing (MicrobesNG, Birmingham, UK). LF82-sGFP2 and LF82 Δ*LF82_314*-mScarlet-I were constructed using pMRE-Tn7-132 and pMRE-Tn7-135 respectively, as in [81]. See Table 2 for a list of plasmids used in this study, and Table 3 for a list of primers.

**Table 2.**

| Plasmid | Description |
| --- | --- |
| pBAD18 | Expression plasmid; Kan<sup>R</sup> |
| pBAD33 | Expression plasmid; Cm<sup>R</sup> |
| pBAD-*LF82_314*-mEmerald | Expression plasmid for LF82_p314-mEmerald fusion protein; Cm<sup>R</sup> |
| pCas9-cr4 | Cas9- expressing plasmid for no-SCAR deletion strategy; Tet<sup>R</sup> |
| pCMV-3xFLAG-*LF82_314* | Plasmid for expression of FLAG-tagged LF82_p314 in mammalian cells; Amp<sup>R</sup> |
| pCMV-mEmerald-4GS | Plasmid for expression of mEmerald-tagged proteins in mammalian cells; Amp<sup>R</sup> |
| pCMV-mEmerald-*LF82_314* | Plasmid for expression of mEmerald-tagged LF82_p314 in mammalian cells; Amp<sup>R</sup> |
| pKD-sgRNA-p314 | λ-Red recombinase- and *LF82_314*-targeting sgRNA-expressing plasmid for no-SCAR deletion strategy; Spc<sup>R</sup> |
| p*LF82_314* | Complementation plasmid for LF82 $\Delta LF82\_314$ encoding an LF82_p314-3xFLAG protein on pBAD18; Kan<sup>R</sup> |
| pMRE-Tn7-132 | Conjugative suicide plasmid for transposon cloning of sGFP2 construct into chromosomal sites; Amp<sup>R</sup>, Cm<sup>R</sup> |
| pMRE-Tn7-135 | Conjugative suicide plasmid for transposon cloning of mScarlet-I construct into chromosomal sites; Amp<sup>R</sup>, Cm<sup>R</sup> |

Plasmids used in this study. Amp<sup>R</sup> = Ampicillin resistant; Cm<sup>R</sup> = Chloramphenicol resistant; Kan<sup>R</sup> = Kanamycin resistant; Spc<sup>R</sup> = Spectinomycin resistant; Tet<sup>R</sup> = Tetracycline resistant.

**Table 3.**
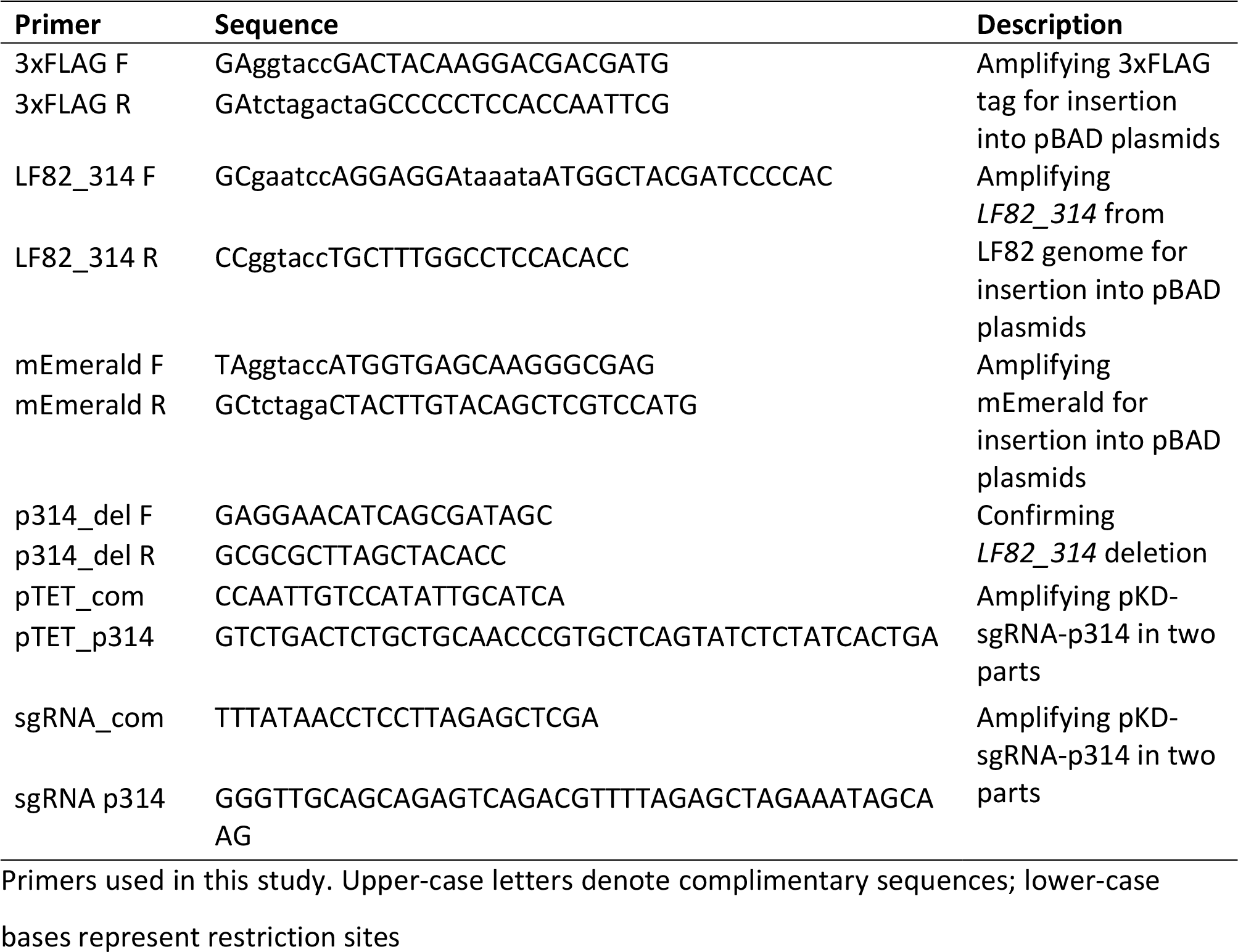

### Light microscopy

Light microscopy was conducted using an Olympus BX51 microscope at appropriate magnifications, using µManager software [82].

### Electron microscopy

EM images were captured using a Tecnai T12 BioTwin Transmission Electron Microscope at an accelerating voltage of 100 kV. Images were captured using a Megaview III Soft Imaging System (SIS) camera.

### Biofilm assays

Crystal Violet (CV) biofilm assays were adapted from [83]. Overnight bacterial cultures were diluted 1:100 in LB, and 100 µl diluted culture was inoculated into each well of a 96-well microtitre plate before static incubation at 37°C. For *trans*-complementation assays, diluted cultures were mixed in specified ratios before inoculation. At appropriate intervals, planktonic bacteria were removed from the plate, and biofilms were washed three times with phosphate buffered saline (PBS). Washed biofilms were stained with 0.1% CV dissolved in water. CV was removed, and stained biofilms were washed four times with PBS, before the plates were dried in a laminar flow cabinet. Dry stain was solubilised in 30% glacial acetic acid and moved to a clean 96-well plate. The OD_550_ of solubilised CV was read using an automated plate reader. Each experiment contained 3-4 technical replicates of 4 biological replicates.

### Biofilm microscopy

Biofilms were grown for microscopy on acid-washed coverslips. Coverslips were placed in 12-well plates, which were then inoculated with 500 µl bacterial culture diluted as above. Inoculated plates were inclined at a 45° angle to ensure the air-liquid interface bisected the coverslip, and were incubated at 37°C for 16 h. At 16 h, culture media was aspirated, and biofilms were fixed in 4% formaldehyde in PBS for 1 h. Fixed biofilms were washed three times with PBS, and coverslips were mounted on slides in a 90% glycerol mounting medium with 0.1% DABCO (Sigma), before imaging at 40× and 100× magnification.

### Motility assays

Motility was assessed using established soft agar protocols. 5µl of saturated overnight culture was inoculated into the centre of soft LB agar plates, solidified with either 0.15% (swimming) or 0.25% (swarming) agar (Sigma) supplemented with 0.4% glucose. Plates were incubated at 37°C. At appropriate intervals, the maximum diameter of the resulting bacterial cloud or swarm was determined, and plates were imaged using a handheld camera.

### Chemotaxis assays

A chemotaxis assay was modified from [84]. 75mm Haemocrit capillary tubes (Hawksley & Sons Ltd, catalogue no. 01604-00) were sealed at one end in a Bunsen flame, before being passed quickly through the flame several times to heat the glass. Heated capillaries were immediately placed open-end down into LB with or without 0.4% (w/v) glucose, and left to draw in media for 15 minutes. Overnight cultures were diluted 1:100 in fresh LB, and inoculated into the wells of a 96-well plate. Media-loaded capillaries were placed into inoculated wells, and the plate was incubated in a laminar flow cabinet at room temperature for 1 h. To recover bacteria, the outside of capillaries were washed with water, and the sealed ends were broken over tubes containing fresh LB to catch escaping culture. Remaining culture was removed by pipetting. Recovered bacteria were then plated at appropriate dilutions for colony forming unit (CFU) enumeration.

### Flagella staining

To prepare bacteria for light microscopy, overnight cultures were diluted 1:33 in fresh LB and incubated at 37°C with agitation for 3 hours, before being spread onto glass slides and stained as in [65]. Stained bacteria were then mounted in immersion oil under a cover slip, sealed with nail varnish, and imaged at 100× magnification. TEM samples were prepared in a protocol modified from [85]. Bacterial cultures were prepared as above, before being absorbed onto carbon-coated copper grids (EM Resolutions) for 10 minutes. Excess fluid was blotted away, and bacteria were fixed in 3% glutaraldehyde in 0.1 M sodium cacodylate buffer for 2 minutes. Fixed samples were washed three times with 0.1 M sodium cacodylate buffer for 10 seconds. Samples were then stained using 2% phosphotungstic acid before imaging at 6000× magnification as above.

### Surface staining

To stain the outer surface of LF82, we employed an amine-reactive dye conjugation, modified from [86]. An overnight culture of LF82 p*LF82_314-mEmerald* was sub-cultured as for flagella staining, and LF82_p314 expression was induced with 0.1% arabinose. At appropriate intervals, 1 ml culture was spun at 1200 × *g* for 10 minutes in a 15ml round-bottom tube, washed twice in 1 ml PBS, and resuspended in 100 µl PBS using minimal agitation. Cells were stained at 37°C using 300 µg/ml CF-633 succinimidyl ester (Sigma) for 30 minutes with agitation at 100 rpm, then washed in PBS. Finally, bacteria were fixed in 4% formaldehyde in PBS for 20 minutes, washed, and mounted on 2% agarose pads for microscopy. Images were captured using a 100× lens.

### *C. elegans* kill assays

A *C. elegans* infection model was modified from [66]. To preclude data being confounded by progeny, *C. elegans* SS104 cultures were synchronised as in [79], and maintained at 25°C to ensure development of sterile adult worms. To prepare kill plates, 100 µl 10× concentrated overnight cultures was spread onto NGM agar plates with appropriate antibiotics, and incubated at 37°C overnight. 30 synchronised L4 or young adult worms were transferred to the prepared kill plates, and incubated at 25°C for up to 14 days. Plates were scored every 24 hours for death, and dead worms were removed. To prevent contamination, worms were transferred to freshly prepared kill plates every 4 days.

### *C. elegans* colonisation assays

*C. elegans* colonisation assays were modified from [66]. Colonisation plates were prepared as above, and 50 synchronised L4 or young adult worms were transferred to each plate, before being incubated at 25°C. To assay stable gut colonisation, 10 worms were transferred to freshly prepared OP50 lawns daily, and incubated at 25°C for 1 h to “wash” transient bacteria from the gut and worm exterior. Worms were then picked and suspended in 1ml M9 buffer, before being washed three times by pelleting at 1200 × *g* for 1 minute before removal of 750 µL buffer, which was replaced with fresh M9 buffer. To determine external bacterial numbers following washing, a sample from the final wash was plated at appropriate dilutions on selective agar. To release gut contents, the worms were homogenised by vortexing with approximately 400 mg sterile 1mm diameter glass beads (BioSpec Products Inc., catalogue no. 11079110) in 1% Triton-X (Sigma) in M9 buffer for 2 minutes, before plating the homogenate at appropriate dilutions on selective agar. CFU per worm gut was defined as:

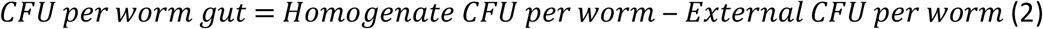

Colonisation experiments were conducted at least twice, with three separate biological replicates per experiment.

### *C. elegans* competition assays

Plates for competition assays were prepared as above, using 1:1 ratios of LF82-Kan^R^:LF82 Δ*LF82_314*-Cm^R^ or LF82-Cm^R^:LF82 Δ*LF82_314*-Kan^R^. Plates with an input ratio substantially different from 1 were discarded. CFU per worm gut was assessed as above, and a competitive index (CI) was calculated. CI was defined as:

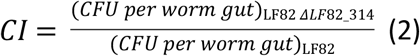

Competition assays were conducted three times, with three separate biological replicates of each mix per experiment.

To image competition assays, infected worms were “washed” on OP50 lawns as above, and immobilised in a 0.1% NaN_3_ solution on a 2% agarose 0.05% NaN_3_ pad on a glass slide, which was sealed under a cover slip. Slides were imaged at 10× magnification.

### Data analysis

Statistical analyses were conducted in GraphPad Prism 7. Error bars in graphs represent standard deviation, unless stated otherwise. Light microscopy images were analysed and processed using FIJI. Fluorescence co-localisation data analysis and Pearson correlation analysis was conducted in CellProfiler. Raw *r* values were converted to z’ values by Fisher’s Z-Transformation and used to calculate mean correlations and 95% CIs, before transformation back to Pearson’s *r* values for interpretation. Cross-sectional intensity measurements were taken using ImageJ.

### Bioinformatics

Protein and nucleotide sequences were retrieved from NCBI [87]. 16S rRNA sequences were harvested from SILVA [88]. Protein homology searches were conducted using BLASTp and JACKHMMER [89,90]. Nucleotide sequences for phylogenetic analysis were retrieved by discontiguous megablast [91]. Multiple sequence alignments were generated using MAAFT [92–94], before submission to PhyML [95] for automated tree generation. Trees were visualised in PRESTO (Phylogenetic tReE viSualisaTiOn, available at http://www.atgc-montpellier.fr/presto/).

## Acknowledgements

The authors thank the Nanoscale and Microscale Research Centre (nmRC, University of Nottingham, UK) for providing access to instrumentation. We would also like to thank Eric J G Pollitt (University of Sheffield, UK) for help in preparing the manuscript.

## Supporting information

**S1 Fig. *LF82_314* self-assembles into large filaments** When expressed heterologously in HeLa cells, *LF82_314* forms large filaments. Filament formation is independent of the tag used for visualisation, suggesting this phenotype is not a tag-dependent artefact. See S3 File Supplementary methods for protocol.

**S2 Fig. *LF82_314* homologue distribution does not mirror true phylogenies** The distribution of *LF82_314* homologues represented in the *LF82_314* ML phylogenetic tree (A) differs significantly from the 16S rRNA phylogenies (B). For example, in (A), *E. coli* LF82 and NRG 857c cluster with *Shigella boydii* Sb227 and ATCC 9210 in a separate clade to other *E. coli* isolates; however, in (B), LF82, NRG 857c, Sb227, and ATCC 9210 cluster as expected with *E. coli* in a distinct clade. Similarly, (A) suggests a loose relationship exists between *Salmonella* strains, whereas in (B), a distinct phylogenetic group is generated. These data suggest *LF82_314* may be horizontally inherited. Strains are coloured by genus. Scale bars = substitutions per site.

**S1 File *LF82_314* is co-inherited with the general secretion pathway**

**S2 File *LF82_314* homologues from blastn discontiguous megablast**

**S3 File Supplementary methods**

**S1 Table p-values for Fig. 1**

